# High-quality *Arabidopsis thaliana* Genome Assembly with Nanopore and HiFi Long Reads

**DOI:** 10.1101/2021.06.08.447650

**Authors:** Bo Wang, Xiaofei Yang, Yanyan Jia, Yu Xu, Peng Jia, Ningxin Dang, Songbo Wang, Tun Xu, Xixi Zhao, Shenghan Gao, Quanbin Dong, Kai Ye

**Author notes:** Corresponding authors. (Ye K), (Yang X).

## Abstract

*Arabidopsis thaliana* is an important and long-established model species for plant molecular biology, genetics, epigenetics, and genomics. However, the latest version of reference genome still contains significant number of missing segments. Here, we report a high-quality and almost complete Col-0 genome assembly with two gaps (Col-XJTU) using combination of Oxford Nanopore Technology ultra-long reads, PacBio high-fidelity long reads, and Hi-C data. The total genome assembly size is 133,725,193 bp, introducing 14.6 Mb of novel sequences compared to the TAIR10.1 reference genome. All five chromosomes of Col-XJTU assembly are highly accurate with consensus quality (QV) scores > 60 (ranging from 62 to 68), which are higher than those of TAIR10.1 reference (QV scores ranging from 45 to 52). We have completely resolved chromosome (Chr) 3 and Chr5 in a telomere-to-telomere manner. Chr4 has been completely resolved except the nucleolar organizing regions, which comprise long repetitive DNA fragments. The Chr1 centromere (CEN1), reportedly around 9 Mb in length, is particularly challenging to assemble due to the presence of tens of thousands of CEN180 satellite repeats. Using the cutting-edge sequencing data and novel computational approaches, we assembled about 4 Mb of sequence for CEN1 and a 3.5-Mb-long CEN2. We investigated the structure and epigenetics of centromeres. We detected four clusters of CEN180 monomers, and found that the centromere-specific histone H3-like protein (CENH3) exhibits a strong preference for CEN180 cluster 3. Moreover, we observed hypomethylation patterns in CENH3-enriched regions. We believe that this high-quality genome assembly, Col-XJTU, would serve as a valuable reference to better understand the global pattern of centromeric polymorphisms, as well as genetic and epigenetic features in plants.

## Introduction

The *Arabidopsis thaliana* Col-0 genome sequence was published in 2000 [1], and after decades of work, this reference genome has become the “gold standard” for *A. thaliana*. However, centromeres, telomeres, and nucleolar organizing regions (NORs) have been either misassembled or not even been sequenced yet due to the enrichment of highly repetitive elements in these regions [2,3]. Long-read sequencing technologies, such as Oxford Nanopore Technology (ONT) sequencing and Pacific Biosciences (PacBio) single molecule real-time (SMRT) sequencing, generate single molecular reads longer than 10 kb, which exceeds the length of most simple repeats in many genomes, making it possible to achieve highly contiguous genome assemblies [4]. Highly repetitive regions, *e*.*g*., centromere or telomere regions, however, remain mostly unassembled due to the limitations in read length and the error rate associated with sequencing of long reads. Although ONT sequencing has overcome read length limitation and can generate ultra-long reads (longest > 4 Mb) (https://nanoporetech.com/products/promethion), the associated 5%□15% per base error rate [5] leads to misassembly or inaccurate assemblies. Naish et al. [6] used ONT-generated ultra-long reads to produce a highly contiguous *A. thaliana* Col-0 genome, but the consensus quality (QV) scores of all five chromosomes, ranging from 41 to 43, were lower than those of the reference TAIR10.1 [6]. High-fidelity (HiFi) data generated from a circular consensus sequencing [7] is a promising strategy for repeat characterization and centromeric satellite assembly. The combination of ONT long reads and HiFi reads has been demonstrated to overcome the issues of sequencing centromere and telomere regions in the human genome and generate the telomere-to-telomere (T2T) assembly of human chromosome (Chr) X [8] and Chr8 [9].

Centromeres mainly consist of satellite DNAs and long terminal repeat (LTR) retrotransposons [10] that attract microtubule attachment and play an important role in maintaining the integrity of chromosomes during cell division [11]. In plant species, centromeric satellite DNA repeats range from 150 to 180 bp in size [12]. It has been reported that *A. thaliana* centromeres contain megabase-sized islands of 178-bp tandem satellite DNA repeats (CEN180) [13] that bind to centromere-specific histone H3-like protein (CENH3) [14,15]. Unfortunately, centromere sequences are largely absent from previously generated *A. thaliana* reference genome assemblies [15], hindering the investigation of CEN180 distribution and its genetic and epigenetic impact on the five chromosomes.

To obtain T2T *A. thaliana* genome assembly, we introduced a bacterial artificial chromosome (BAC)-anchor replacement strategy to our assembly pipeline and generated the Col-XJTU genome assembly of *A. thaliana*. We completely resolved the centromeres of Chr3, Chr4, Chr5, and partially resolved centromeres of Chr1 and Chr2. Col-XJTU assembly of *A. thaliana* genome was found to be highly accurate with a QV score greater than 60, which is obviously higher than that of TAIR10.1 and another recently deposited assembly [6]. Due to the unprecedented high quality of the Col-XJTU genome assembly, we were able to observe intriguing genetic and epigenetic patterns in the five centromere regions.

## Results

### Assembly of a high-quality genome of *A. thaliana*

We assembled ONT long reads using NextDenovo v. 2.0, and initially generated 14 contigs (contig N50 = 15.39 Mb) (**Figure 1**A and Figure S1A). Of these, eight contigs contained the *Arabidopsis*-type telomeric repeat unit (CCCTAAA/TTTAGGG) on one end while two contigs had the 45S rDNA units on one end (Figure 1A). Contig 13 (935 kb) and Contig 14 (717 kb) composed of CEN180 sequences were neither ordered nor oriented, and they were removed from the assembly (Figure S1A). We polished the remaining 12 contigs with HiFi data using Nextpolish and scaffolded them using 3D-DNA derived from Hi-C data. Consequently, we obtained five scaffolds with seven gaps located at centromere regions (Figure 1A). To further improve the genome assembly, we assembled HiFi reads using hifiasm [16,17] and identified the centromeric flanking BAC sequences [18-20] on both the five ONT scaffolds and HiFi contig pairs (Figure 1A and Figure S1B, C). We first filled the gaps on centromeres using the BAC-anchor strategy (Figure S1B). To guarantee the highest base-pair accuracy, we replaced the low accuracy ONT-genome-assemblies with the PacBio HiFi contigs and kept the HiFi contigs as long as possible (Figure 1A and Figure S1C). The final genome assembly (contig N50 = 22.25 Mb; scaffold N50 = 26.16 Mb) was named Col-XJTU. The Col-XJTU genome size is 133,725,193 bp (Chr1: 32,659,241 bp; Chr2: 22,560,461 bp; Chr3: 26,161,332 bp; Chr4: 22,250,686 bp; and Chr5: 30,093,473 bp), and the QV scores of all five chromosomes are greater than 60 (ranging from 62 to 68), which are obviously higher than those of the TAIR10.1 reference genome (ranging from 45 to 52) (**Table 1**) and a recently deposited genome (ranging from 41 to 43) [6], suggesting that our Col-XJTU assembly is highly accurate. The completeness evaluation showed a k-mer completeness score of 98.6%, suggesting that Col-XJTU assembly is highly complete as well. The Col-XJTU assembly was composed of 97% HiFi contigs and only 4,098,671-bp-long ONT contigs, which contain highly repetitive elements (Table S1). The heterozygosity of *A. thaliana* Col-XJTU is very low (0.0865%), which was estimated using GenomeScope v. 1.0 [21] from the k-mer 17 histogram computed by Jellyfish v. 2.3.0 [22]. The base accuracy and structure correctness of the Col-XJTU assembly was also estimated from sequenced BACs. Firstly, 1465 BACs were aligned to the Col-XJTU assembly via Winnowmap2, and the mapping results calculated using the CIGAR string revealed good agreement with high sequence identity (99.87%). We validated the structure of our assembly using bacValidation, and the Col-XJTU assembly resolves 1427 out of 1465 validation BACs (97.41%), which is higher than BAC resolving rate of humans [23]. In addition, Col-XJTU genome assembly corrected one misassembled region with 1816 bp in length, containing two protein-coding genes, in the TAIR10.1 genome (Figure 1B and Table S2).

**Table 1.** Comparison of genomic features for Col-XJTU and TARI10.1 assemblies.

| **Feature** | **Col-XJTU** | **TAIR10.1** |
| --- | --- | --- |
| Genome size (bp) | 133,725,193 | 119,668,634 |
| GC content | 36.34% | 36.03% |
| Contig N50 (bp) | 22,250,686 | 11,194,537 |
| Scaffold N50 (bp) | 26,161,332 | 23,459,830 |
| QV (accuracy) for Chr1 | 67.78 (99.999983%) | 48.46 (99.998574%) |
| QV (accuracy) for Chr2 | 61.89 (99.999935%) | 52.30 (99.999411%) |
| QV (accuracy) for Chr3 | 66.16 (99.999976%) | 51.27 (99.999254%) |
| QV (accuracy) for Chr4 | 66.73 (99.999979%) | 44.70 (99.996611%) |
| QV (accuracy) for Chr5 | 63.95 (99.999960%) | 48.76 (99.998670%) |
| No. of protein-coding genes | 27,583 | 27,444 |
| Repeat content | 23.87% | 16.23% |
| No. (percentage) of complete BUSCOs | 932 (97.5%) | 931 (97.4%) |
| No. (percentage) of complete and single-copy BUSCOs | 911 (95.3%) | 910 (95.2%) |
| No. (percentage) of complete and duplicated BUSCOs | 21 (2.2%) | 21 (2.2%) |
| No. (percentage) of fragmented BUSCOs | 3 (0.3%) | 3 (0.3%) |
| No. (percentage) of missing BUSCOs | 21 (2.2%) | 22 (2.3%) |
| No. of total BUSCO groups searched | 956 | 956 |

*Note:* QV, consensus quality. Genome completeness was assessed using BUSCO in the “genome” running mode.

**Figure 1.**
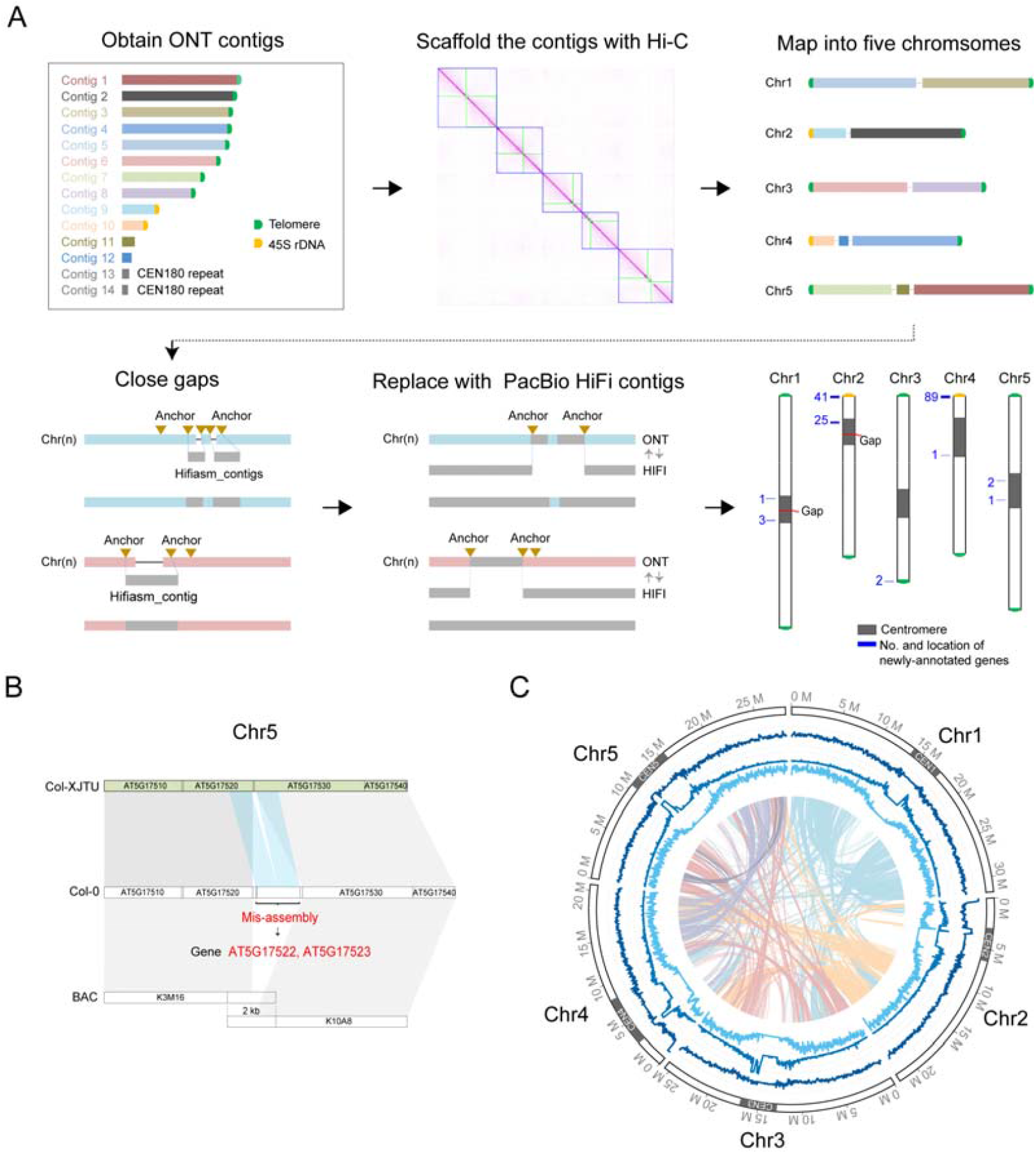
High-quality telomere-to-telomere genome assembly. **A**. ONT ultra-long reads assembly. We obtained 14 contigs, and two of them are composed of CEN180 repeat sequences. Then, the 12 contigs were ordered and oriented to five scaffolds using Hi-C data. After scaffolding, the assembly was represented in five pseudomolecules corresponding to the five chromosomes of the TAIR10.1 assembly. Gaps were filled using HiFi assembly contigs based on BAC anchors. Whole ONT-Hi-C assemblies were replaced with HiFi contigs based on BAC anchors. The gray bars represent the centromere regions, red line represent the gap location; blue number and line indicates the number and locus of new gene annotations. **B**. Col-XJTU genome assembly corrected a misassembly region of TAIR10.1 genome assembly. Grey bands connect corresponding collinear regions. Duplicated segments that were misassembled are connected with bands in blue. **C**. Circos plot of Col-XJTU genome assembly. The tracks from outside to inside: distribution of karyotypes of assembled chromosomes, GC density, density of transposable elements, and gene density calculated in 50-kb windows. Syntenic blocks and different colored lines represent different chromosomes. Chromosome are labeled at the outmost circle with centromeres shown in dark gray. BAC, bacterial artificial chromosome.

The assembly sizes of Col-XJTU centromere 1 (CEN1), CEN2, CEN3, CEN4, and CEN5 were 3.8 Mb, 3.5 Mb, 4.0 Mb, 5.5 Mb, and 4.9 Mb, respectively (Table S3). The sizes of gap-free CEN3, CEN4, and CEN5 were consistent with the physical map-based centromeric sizes [18-20], while the 3.8-Mb-long CEN1 had a gap and was smaller than the estimated size of 9 Mb based on the physical map [20] and the 3.5-Mb-long CEN2 with a gap was assembled 88% of the 4-Mb-long physical map [20]. All five centromeric CEN180 arrays do not contain large structural errors (Figure S2). Upon annotation of the five centromere regions, we found that all five *A. thaliana* centromeres were surrounded by transposon-enriched sequences rather than protein-coding gene-enriched sequences (Figure 1C).

The Col-XJTU assembly (contig N50 = 22.25 Mb) improved the contiguity of the *A. thaliana* genome compared to TAIR10.1 (contig N50: 11.19 Mb) (Table 1) and we had filled 36 gaps apart from two gaps in CEN1 and CEN2 (Table S4). Benchmarking Universal Single-Copy Orthologs (BUSCO) evaluation revealed higher genome completeness of Col-XJTU than that of TAIR10.1 (Table 1). The synteny plot shows that Col-XJTU genome is highly concordant with TAIR10.1 (Figure S3) but with additional completely resolved three centromere regions and partly resolved NORs. Novel sequences (a set of regions not covered by TAIR10.1) equivalent to a total of 14.6 Mb were introduced in the Col-XJTU genome; of these, 94.8% belong to the centromeric regions, with 3.7% of them located in the NORs and telomeres (Table S5). We estimated the QV score of the novel sequences (>10 kb) as 67.43, and the base accuracy is 99.999982%. The assembly sizes of 45S rDNA units in Chr2 and Chr4 were 300,270 and 343,661 bp, respectively. The telomeres of the eight chromosome arms ranged from 1862 to 3563 bp in length (Table S6), which are consistent with the reported lengths [24]. The read depths of these telomeres did not differ obviously compared to the average coverage of the genome (Table S6). Moreover, no telomeric motifs were found in the unmapped HiFi reads, probably indicating completely resolved telomeres. The repeat content of Col-XJTU genome (24%) is much higher than that of the current reference genome (16%) (Table 1), largely due to higher number of LTR elements assembled and annotated in Col-XJTU genome (Table S7).

A total of 27,418 protein-coding genes (99.9%) were lifted-over from TAIR10.1 (27,444) using Liftoff (Table 1). We then masked repeat elements and annotated protein-coding genes in the novel sequences in Col-XJTU genome. Finally, we obtained 27,583 protein-coding genes in Col-XJTU genome with 165 newly-annotated genes. Of the newly-annotated genes, 41 and 89 genes were located in NORs of Chr2 and Chr4, respectively (Figure S4), while 35 newly-annotated genes were located at centromeres (n = 33) and telomeres (n = 2) (Figure 1A). Only 14 of the 165 newly-annotated genes contain functional domains, whereas the remaining 151 ones have unknown functions (Table S8). Interestingly, 96% of the newly-annotated genes were found to be actively transcribed across different tissues (Table S9), especially in leaves (Figure S5). The highly expressed leaf-specific novel genes encode protein domains such as ATP synthase subunit C and NADH dehydrogenase (Table S8), indicating that these genes may be involved in photosynthesis.

### Global view of centromere architecture

Previously, the centromere composition of *A. thaliana* was estimated using physical mapping and cytogenetic assays; however, such estimation resulted in the generation of incorrectly annotated and unknown regions, such as 5S rDNA and CEN180 repeat regions [1]. The complete assembly of CEN3, CEN4, and CEN5 in this study revealed ∼0.5 kb-long repeats in the 5S rDNA array regions (**Figure 2**), which is consistent with the findings of previous studies that used fluorescence *in situ* hybridization and physical mapping [25,26]. The 5S rDNA regions in CEN4 and CEN5 exhibited high similarity with 95% sequence identity. However, this region in CEN3 is interrupted by LTRs, resulting in a low sequence identity. All 5S rDNA regions present GC-rich and hypermethylation patterns (Figure 2). We detected 3666 5S rDNA monomers, which approximately doubles the previously reported amount of ∼2000 5S rDNA gene copies in the Col-0 genome [27]. The 5S rDNA arrays were divided into four clusters (Figure S6A), wherein the 5S rDNA from Chr4 as well as Chr5 formed independent clusters labeled as 5S cluster 1 and 2, respectively (Figure 2). The 5S rDNA sequences on Chr3 were divided into two 5S clusters (Figure 2), which contain obviously more polymorphic sites than that of Chr4 and Chr5 (Figure S7).

**Figure 2.**
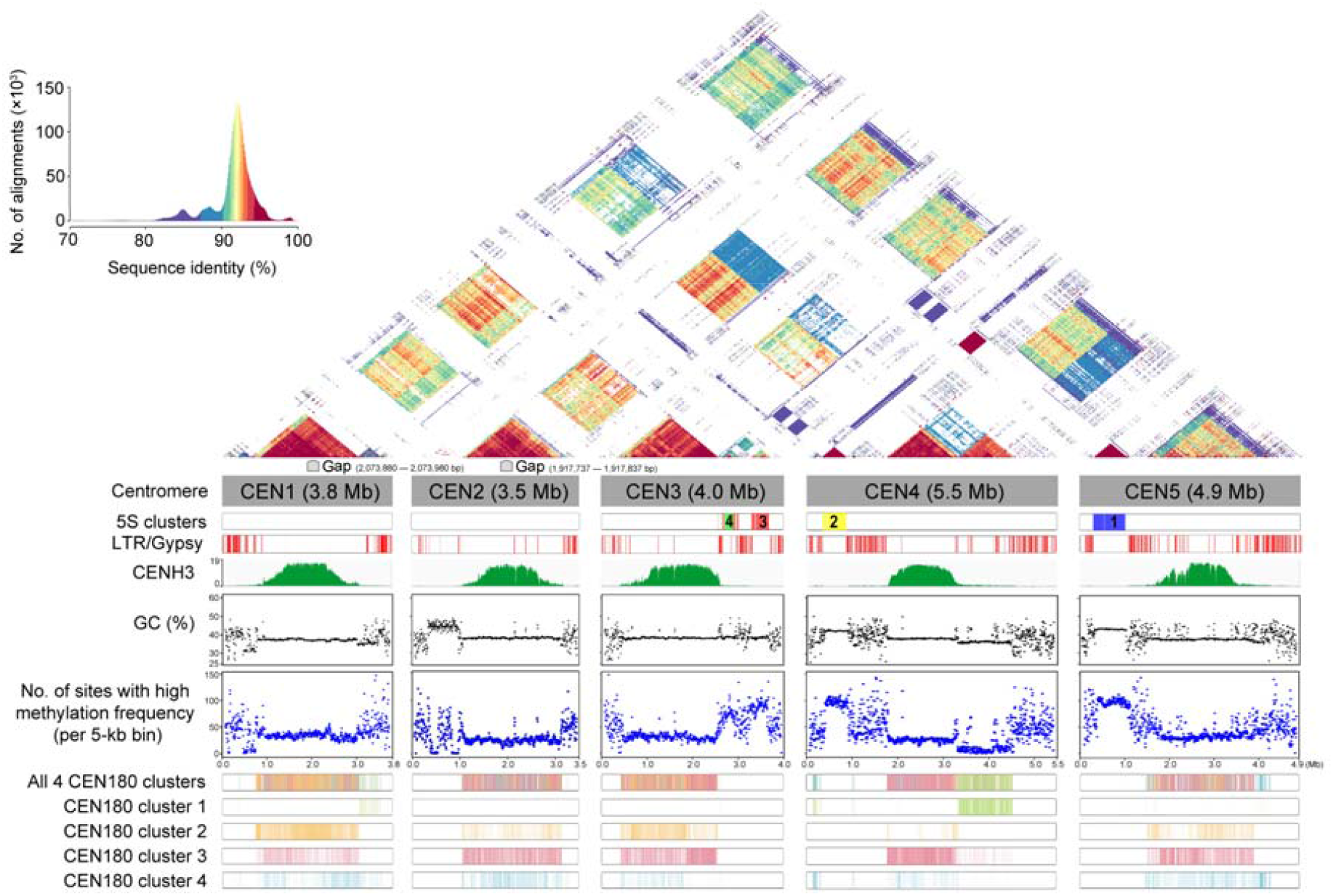
Structure and epigenetic map of the five chromosome centromeric regions. Global view of the patterns of 5S rDNA, LTR/Gypsy, GC content, CENH3 ChIP-seq binding map, and CpG methylation frequency (> 0.8) of the five centromeric regions. The four CEN180 clusters and four 5S clusters are shown as color bars. 5S clusters 1–4 are shown in blue, yellow, red, and green, respectively. LTR, long terminal repeat; CENH3, centromere-specific histone H3-like protein.

We observed that CEN1, CEN2, CEN3, and CEN4 contained highly similar CEN180 arrays (Figure 2), but the reduced internal similarity in CEN5 was likely due to the disruption of LTR/Gypsy elements (Figure 2). We found one CEN180 array in CEN1, CEN2, CEN3, and CEN5 but two distinct CEN180 arrays in CEN4. Except for the downstream one in CEN4, all others CEN180 arrays showed higher than 90% sequence identity either with inter- or intra-chromosomal regions (Figure 2). The downstream CEN180 array in CEN4 showed a higher internal sequence identity (> 90%) and a lower external sequence identity (< 90%) than the other CEN180 arrays (Figure 2). Moreover, the downstream CEN180 array in CEN4 showed lower GC content and methylation frequency than other CEN180 arrays (Figure 2). We performed LASTZ searches for tandem repeats to construct the CEN180 satellite library and identified 60,563 CEN180 monomers in the five centromeres. The phylogenetic clustering analysis revealed that four distinct CEN180 clusters with single-nucleotide variants and small indels (Figure S6B and Figure S8). Almost all the downstream CEN180 arrays of CEN4 belong to CEN180 cluster 1 (Figure 2 and Figure S9), while the upstream CEN180 arrays of CEN4 belong to the remaining three CEN180 clusters.

A functional region of centromere is defined by the binding of epigenetic modifications with CENH3 [28,29]. We observed that CENH3 is obviously enriched in the interior of the centromere but depleted at the LTR region (Figure 2). The five centromeres showed higher DNA methylation than pericentromeres (Figure S10); however, the CEN180 arrays presented hypomethylation patterns (Figure 2 and Figure S10). Interestingly, we found that the CENH3-binding signal exhibits a strong preference for CEN180 cluster 3 on all five centromeres (Figure S11). Such preference is observed in CEN180 cluster 1 in CEN4 and other four centromeres (Figure S11). The CENH3 signal enrichment presented the opposite tendency with the methylation frequency in 60% CEN180 clusters of the five centromeres (Figure 2 and Figure S12).

## Discussion

Traditionally, long-read sequencing technologies commonly suffer from high error rates [30]. However, the recently developed HiFi reads by PacBio have both the advantages of long read lengths and low error rates, enabling the assembly of complex and highly repetitive regions in the new era of T2T genomics [31,32]. HiFi reads were used to assemble the T2T sequence of human ChrX and Chr8 [8,9], aiding in the completion of the human genome [33]. Recently, two complete rice reference genomes have also been assembled using HiFi reads [32].

The size of *A. thaliana* centromeres is 2–5 folds larger than that of the rice centromeres (0.6–1.8 Mb) [32], and hence, a sophisticated approach is required to complete the assembly of the *A. thaliana* centromeres. We combined the dual long-read platforms of ONT ultra-long and PacBio HiFi to produce the high-quality *A. thaliana* Col-XJTU genome with only two gaps in CEN1 and CEN2. The CEN1 is 3.8 Mb long and is smaller than the 9-Mb region estimated by physical mapping [20]. We assembled a 3.5-Mb-long sequence (88% of the physical map [20]) of CEN2 using hifiasm. Recently, a version of *A. thaliana* genome was deposited with ∼5-Mb-long CEN1 sequence, which is still smaller than the physical map size [20], indicating the difficulty in assembling long centromere regions even with long-read technologies [6]. We are optimizing a singly unique nucleotide *k*-mers (SUNKs) assembly method [9] for plant genomes, aiming to eventually produce the completely resolved long centromere regions.

Diverse methylation patterns have been observed in the centromere sequences of two human chromosomes upon completion of the human genome [8,9]. The centromeres of Chr8 and ChrX in the human genome contain a hypomethylation pocket, wherein the centromeric histone CENP-A for kinetochore binding is located [8,9,34,35]. This phenomenon has also been observed experimentally in *A. thaliana* [36]. Our high-quality centromere assembly of *A. thaliana* reveals that the CEN180 arrays enriched with CENH3 occupancy are hypomethylated compared to the pericentromeric regions. Although the primary function of centromeres is conserved between animal and plant kingdoms, the centromeric repeat monomers are highly variable in terms of sequence composition and length, and little sequence conservation is observed between species [37]. Extensive experimental evidence has confirmed that convergent evolution of centromere structure, rather than the sequence composition, is the key to maintaining the function of centromeres [38]. Furthermore, we have observed clusters with irregular patterns of methylation and CENH3 binding, indicating that centromeres may contain regions with unknown functions or still-evolving components. We would need to complete the assembly of centromere sequence for more related species to gain insight into the evolution of centromere structure and function.

In conclusion, our novel assembly strategy involving the combination of ONT long reads and HiFi reads leads to the assembly of a high-quality genome of the model plant *A. thaliana*. This genome will serve as the foundation for further understanding molecular biology, genetics, epigenetics, and genome architecture in plants.

## Materials and methods

### Plant growth condition and data sources

The *A. thaliana* accession Col-0 was obtained from the Shandong Agricultural University as a gift. The *A. thaliana* seeds were placed in a potting soil and then maintained in a growth chamber at 22 °C with a 16 h light/8 h dark photoperiod and a light intensity of 100–120 µmol m^-2^ s^-1^. Young true leaves taken from 4-week-old healthy seedlings were used for sequencing.

The Illumina short read data of our Col-0, public wild type Col-0 (JGI Project ID: 1119135), and another accession AT1741 (SRA: ERR2173372) were mapped to the reference genome TAIR10.1 (RefSeq: GCF_000001735.4). We used a series of software packages, including bwa v. 0.7.17–r1188 [39], biobambam v. 2.0.87, samtools v. 1.9 [40], varscan v. 2.4.4 [41], bcftools v.1.9 (https://samtools.github.io/bcftools), and tabix v. 1.9 [42], for single nucleotide polymorphism (SNP) calling. The SNP calling results indicated that Col-XJTU was highly similar to the public wild type Col-0 (Figure S13).

### Genomic DNA sample preparation

DNA was extracted using the Qiagen® Genomic DNA Kit following the manufacturer guidelines (Cat#13323, Qiagen, Valencia, CA). Quality and quantity of total DNA were evaluated using a NanoDrop™One UV-Vis spectrophotometer (Thermo Fisher Scientific, Waltham, MA) and Qubit® 3.0 Fluorometer (Invitrogen life Technologies, Carlsbad, CA), respectively. The Blue Pippin system (Sage Science, Beverly, MA) was used to retrieve large DNA fragments by gel cutting.

### Oxford Nanopore PromethION library preparation and sequencing

For the ultra-long Nanopore library, approximately 8–10 µg of genomic DNA was selected (> 50 kb) with the SageHLS HMW library system (Sage Science), and then processed using the Ligation sequencing 1D kit (SQK-LSK109, Oxford Nanopore Technologies, Oxford, UK) according the manufacturer’s instructions. DNA libraries (approximately 800 ng) were constructed and sequenced on the Promethion (Oxford Nanopore Technologies) at the Genome Center of Grandomics (Wuhan, China). A total of 56.54 Gb of ONT long reads with ∼388 × coverage were generated including ∼177 × coverage of ultra-long (> 50 kb) reads. The N50 of ONT long reads is 46,452 bp, and the longest reads were 495,032 bp.

### ONT long reads assembly and correction

The long-read assembler NextDenovo v. 2.0 (https://github.com/Nextomics/NextDenovo) was used to assemble the ONT long reads with parameters: ‘read_cutoff = 5k’ and ‘seed_cutoff = 108,967’. Nextpolish v. 1.3.0 [43] with parameters ‘hifi_options -min_read_len 10k -max_read_len 45k -max_depth 150’ was used to polish the contigs assembled by ONT long reads.

### HiFi sequencing and assembly

SMRTbell libraries were constructed according to standard protocol of PacBio using 15 kb preparation solutions (Pacific Biosciences, CA). The main steps for library preparation include: (1) genomic DNA shearing; (2) DNA damage repair, end repair, and A-tailing; (3) ligation with hairpin adapters from the SMRTbell Express Template Prep Kit 2.0 (Pacific Biosciences); (4) nuclease treatment of SMRTbell library with SMRTbell Enzyme Cleanup Kit; and (5) size selection and binding to polymerase. The 15 µg genomic DNA sample was sheared by gTUBEs. Single-strand overhangs were then removed, and DNA fragments were damage repaired, end repaired, and A-tailed. Then, the fragments were ligated with the hairpin adapters for PacBio sequencing. The library was treated with the nuclease provided in the SMRTbell Enzyme Cleanup Kit and purified by AMPure PB Beads. Target fragments were screened by BluePippin (Sage Science). The SMRTbell library was then purified by AMPure PB beads, and Agilent 2100 Bioanalyzer (Agilent Technologies, Palo Alto, CA) was used to detect the size of the library fragments. Sequencing was performed on a PacBio Sequel II instrument with Sequencing Primer V2 and Sequel II Binding Kit 2.0 at the Genome Center of Grandomics. A total of 22.90 Gb of HiFi reads with ∼157 × coverage were generated, and N50 of the reads was 15,424 bp. HiFi reads were assembled using hifiasm v. 0.14-r312 [17] with default parameters, and the gfatools (https://github.com/lh3/gfatools) was used to convert sequence graphs in the GFA to FASTA format.

### Hi-C sequencing and scaffolding

Hi-C library was prepared from cross-linked chromatins of plant cells using a standard Hi-C protocol; the library was then sequenced using Illumina NovaSeq 6000. A total of 21.14 Gb of Hi-C reads with ∼158 × coverage were generated. The Hi-C sequencing data were used to anchor all contigs using Juicer v. 1.5 [44], followed by a 3D-DNA scaffolding pipeline [45]. Scaffolds were then manually checked and refined with Juicebox v. 1.11.08 [46].

### Replacing ONT-HiC-assemblies with HiFi-contigs

We introduced a BAC-anchor strategy to fill the remaining gaps in ONT-HiC assemblies. Briefly, for each gap, we first identified two BAC sequences flanking the gap locus that were aligned concordantly (identity > 99.9%) to both the ONT-HiC assembly and a HiFi contig, and then replaced the gap containing contigs with corresponding HiFi contigs. We used the same method to polish ONT-HiC assemblies with HiFi contigs. The BAC sequences we used as anchors are list in Figure S1 and Table S10.

### Genome comparisons

Two genome assemblies were aligned against each other using nucmer v. 4.0.0 (-c 100 -b 500 -l 50) [47], and the output delta file was filtered using a delta-filter (-i 95 -l 50). The alignment regions between two genomes were extracted using show-coords (-c -d -l -I 95 -L 10,000), and the novel region of our genome was extracted using ‘complement’ in BEDTools v. 2.30.0 [48]. The synteny relationships among the five chromosomes were estimated using by BLASTN v. 2.9.0 with ‘all vs.all’ strategy and visualized using Circos v. 0.69-8 [49]. Genomic alignment dot plot between Col-XJTU and TAIR10.1 assemblies was generated using D-GENIES [50]. QV and completeness scores were estimated using Merqury [51] from Illumina sequencing data generated on the same material in this study. The assembly accuracy for five chromosomes was estimated from QV as follows: 100 − (10^(QV/−10)^) × 100 = accuracy percentage [9]. To assess genome completeness, we also applied BUSCO v. 3.0.2 analysis using the plant early release database v. 1.1b [52]. Pairwise sequence identity heat maps of five centromere were calculated and visualized using the following aln_plot (https://github.com/mrvollger/aln_plot) command: bash cmds.sh CEN CEN.fa 5000.

### BAC validation

We validated the assemblies using bacValidation (https://github.com/skoren/bacValidation) with default parameters, which recognizes a BAC as ‘resolve’ within the assembly with 99.5% of the BAC length to be aligned to a single contig. BAC libraries were downloaded from European Nucleotide Archive (ENA), and the BACs used to validate five chromosomes are listed in Table S10.

### Assembly validation of CEN180 array

We applied TandemTools [53] to assess the structure of the centromeric CEN180 array. We first aligned ONT reads (> 50 kb) to the Col-XJTU assembly with Winnowmap2 and extracted reads aligning to the centromeric CEN180 array (Chr1: 14,994,091 – 17,146,102; Chr2: 4,274,401 – 6,365,272; Chr3: 13,673,967 – 15,762,202; Chr4_upstream_part: 4,895,149–6,440,779; Chr4_downstream_part: 6,440,780 – 7,708,273; and Chr5: 12,617,763–14,826,408). Then these extracted ONT reads were inputted in tandemquast.py with the parameters ‘-t 96 --nano {ont_reads.fa} -o {out_dir} CEN.fa’.

### Genome annotation

The software Liftoff v. 1.6.1 (-mm2_options = “-a --end-bonus 5 --eqx -N50 -p 0.5”) [54] was used to annotate protein-coding genes of Col-XJTU assembly based on the reference genome. We then used Augustus v. 2.5.5 (--gff3 = on --genemodel = complete --species = arabidopsis) [55] to annotate the novel regions in the Col-XJTU assembly. Transposable elements and 45S rDNA were identified by RepeatMasker v. 4.0.7 (http://www.repeatmasker.org) (-species ‘arabidopsis thaliana’ -s -no_is -cutoff 255 -frag 20000), and 5S rDNA was detected by TideHunter v. 1.4.3 (https://github.com/yangao07/TideHunter) and predicted by rRNAmmer v.1.2 [56].

### Misassembly evaluation

We first used QUAST v. 5.0.2 [57] to assess the structure accuracy of new assemblies. QUAST parameters were set to ‘quast.py <asm> -o quast_results/<asm> -r <reference> --large-min-alignment 20,000 --extensive-mis-size 500,000 --min-identity 90’ according to previous report [23]. Base on QUAST evaluation, we did not detect any misassembly between Col-XJTU and TAIR10.1 genomes at non-centromeric regions. Furthermore, we detected and labeled one potential mis-assembly due to segmental duplications for Chr5 when mapping the protein-coding gene sequences of TAIR10.1 to Col-XJTU using Liftoff. We aligned the BAC sequences (K3M16 and K10A8) to the different regions between TAIR10.1 and Col-XJTU using BLASTN, supporting that Col-XJTU assembly is correct.

### Gene expression analysis

We chose seven tissues for gene expression analysis [58], namely root (SRA: SRR3581356), flower (SRA: SRR3581693), leaf (SRA: SRR3581681), internode (SRA: SRR3581705), seed (SRA: SRR3581706), silique (SRA: SRR3581708), and pedicel (SRA: SRR3581703). A gene expression profile was created using the TopHat v. 2.0.9 (‘-g 1’) and Cufflinks v. 2.2.1 pipeline [59,60]. Fragments per kilobase of transcript per million fragments mapped (FPKM) values of the seven tissues were used to plot a heatmap using TBtools v. 1.068 [61].

### Centromeric satellite DNA and 5S rDNA cluster analysis

LASTZ (http://www.bx.psu.edu/~rsharris/lastz/) [62] with parameters ‘--coverage = 90 --format = general: score, name1, strand1, size1, start1, end1, name2, strand2, identity, length1, align1’ was used to identify CEN180 (query: AAAAGCCTAAGTATTGTTTCCTTGTTAGAAGATACAAAGACAAAGACTCATATGGACTTCGGCTACACCATCAAAGCTTTGAGAAGCAAGAAGAAGCTTGGTTAG TGTTTTGGAGTCAAATATGACTTGATGTCATGTGTATGATTGAGTATAACAACTTAAACCGCAACCGGATCTT) [15] repeats within the complete centromeres. Then, the CEN180 repeats (ranging from165 to 185 bp) were aligned using Clustal Omega v. 1.2.4 [63] with default parameters. Clustering was performed on this alignment using find.best() function with default init.method in R package ‘phyclust’ [64]. We evaluated a range of clusters (K2–K10) and used the Bayesian information criterion (BIC) inflection point approach to choose the optimal K value [64]. The 5S rDNA cluster analysis was performed using the same aforementioned pipeline, and the 5S rDNA repeats were retained from 490 to 510 bp.

### ChIP-seq analysis

The ChIP-seq paired-end reads downloaded from SRA (SRA: SRR4430537 for replicate 1, SRA: SRR4430537 for replicate 2 and SRA: SRR4430541 for control) were mapped to the Col-XJTU assembly using the ‘mem’ algorithm of BWA [39], and the mapping results of two replicates were merged. We carried out peak calling using MACS2 [65] with the parameters ‘-t merged.bam -c control.bam -f BAM --outdir ATChipseq -n ATChipseq -B --nomodel --extsize 165 --keep-dup all’. Mapped read counts of each CEN180 cluster were calculated using ‘multiBamSummary’ in deepTools [66].

### Methylation analysis

Nanopolish v. 0.13.2 with the parameters ‘call-methylation --methylation cpg’ was used to measure the frequency of CpG methylation in raw ONT reads. The ONT reads were aligned to whole-genome assemblies via Winnowmap v. 2.0 [67]. The script ‘calculate_methylation_frequency.py’ provided in the methplotlib package [68] was then used to generate the methylation frequency.

## Data availability

The whole genome sequence data reported in this paper have been deposited in the Genome Warehouse [69] in National Genomics Data Center [70] (BioProject: PRJCA005809), Beijing Institute of Genomics, Chinese Academy of Sciences / China National Center for Bioinformation (GWH: GWHBDNP00000000.1) that is publicly accessible at https://ngdc.cncb.ac.cn/gwh. The genome annotation has been deposited in https://dx.doi.org/10.6084/m9.figshare.14913045. The raw sequencing data for the PacBio HiFi reads, ONT long-reads, Illumina short reads, and Hi-C Illumina reads have been deposited in the Genome Sequence Archive [71] at the National Genomics Data Center, Beijing Institute of Genomics, Chinese Academy of Sciences / China National Center for Bioinformation (GSA: CRA004538), and are publicly accessible at http://bigd.big.ac.cn/gsa.

## Code availability

SNP calling pipeline is available for public use at BioCode (https://ngdc.cncb.ac.cn/biocode/tools/BT007246).

## CRediT author statement

**Bo Wang**: Methodology, Software, Formal analysis, Visualization, Writing - Original draft preparation, Writing - Review and Editing. **Xiaofei Yang**: Methodology, Supervision, Writing - Review and Editing. **Yanyan Jia**: Resources, Methodology. **Yu Xu**: Software, Formal analysis. **Peng Jia**: Software, Formal analysis. **Ningxin Dang**: Methodology, Formal analysis. **Songbo Wang**: Methodology, Formal analysis. **Tun Xu**: Formal analysis. **Xixi Zhao**: Formal analysis. **Shenghan Gao**: Methodology. **Quanbin Dong**: Resources. **Kai Ye**: Conceptualization, Methodology, Supervision, Funding acquisition, Writing - Review & Editing. All authors read and approved the final manuscript.

## Competing interests

The authors have declared no competing interests.

## Acknowledgments

This study was supported by the National Natural Science Foundation of China (Grant Nos. 62172325 and 32070663), National Key R&D Program of China (Grant Nos. 2018YFC0910400 and 2017YFC0907500), China Postdoctoral Science Foundation (Grant No. 2020M673420), the Fundamental Research Funds for the Central Universities, and the World-Class Universities (Disciplines) and the Characteristic Development Guidance Funds for the Central Universities, China.

---

**Figure S1.**
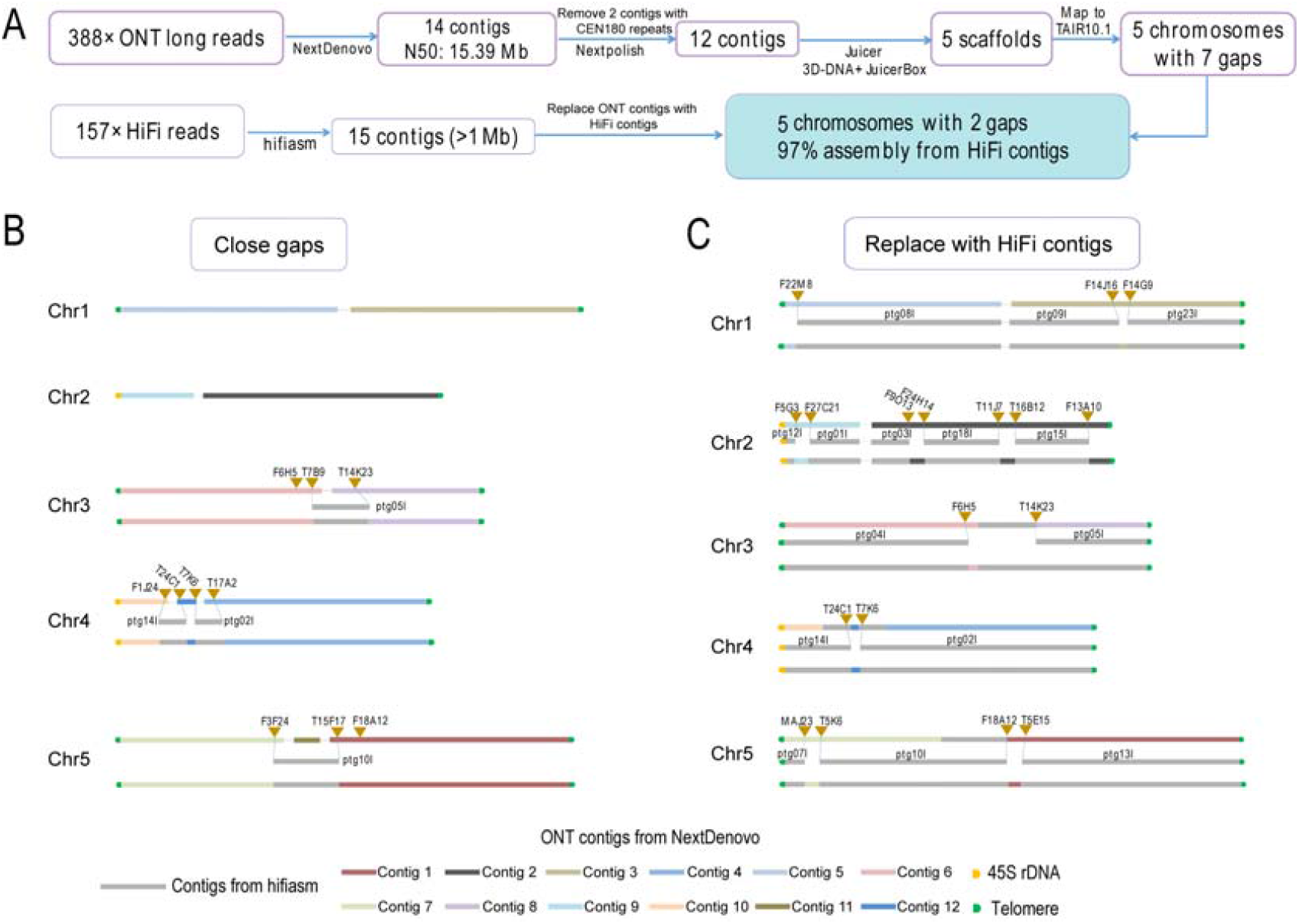
Assembly pipeline and BAC-anchor replacement strategy. **A**. High-quality T2T genome assembly pipeline. **B**. BAC sequences used to close gap. **C**. BAC sequences used to replace ONT contigs with HiFi contigs. All the BACs used share more than 99.9% sequence identity with their target contigs.

**Figure S2.**
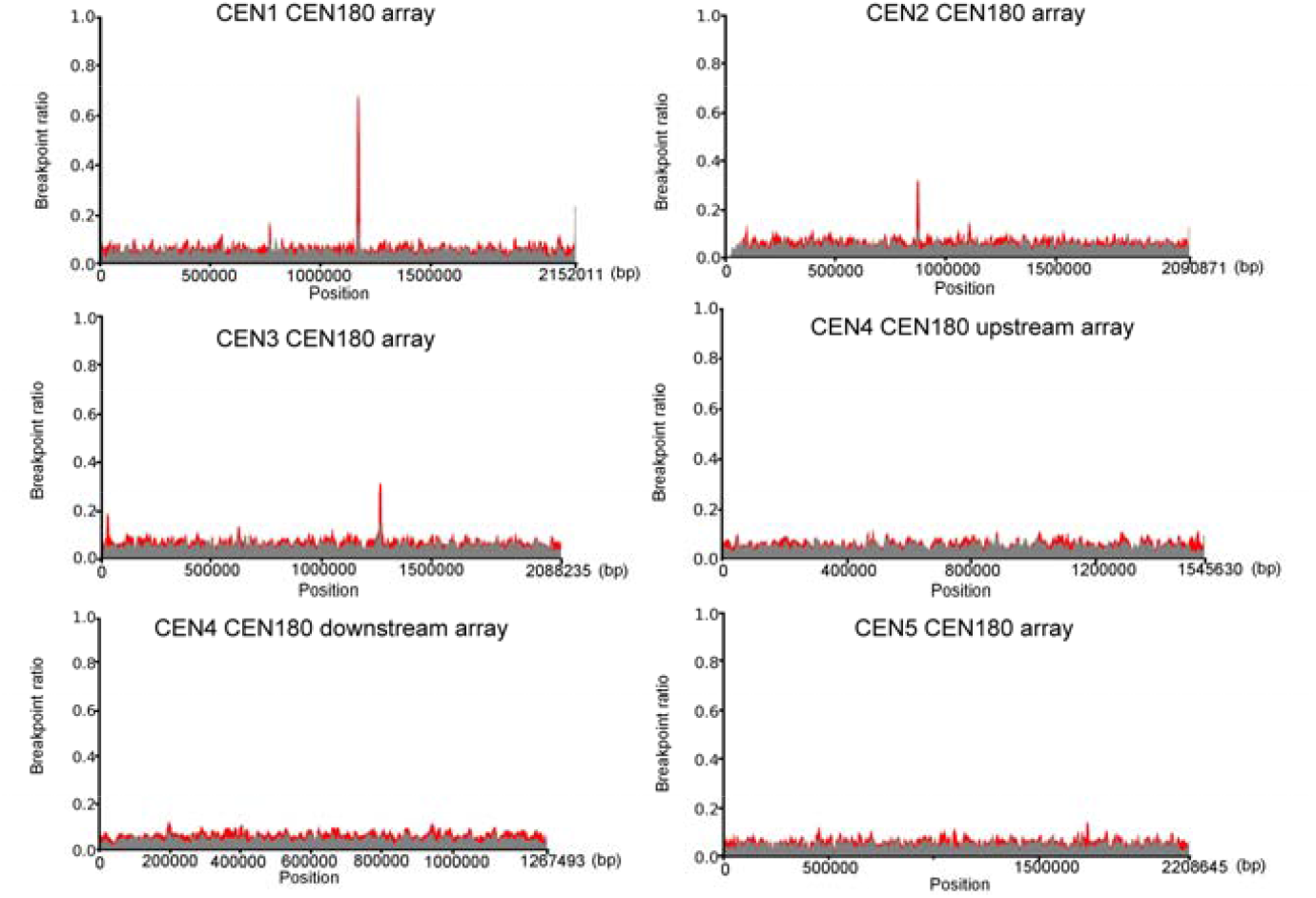
TandemTools validation plots of the five centromeric CEN180 array regions.

**Figure S3.**
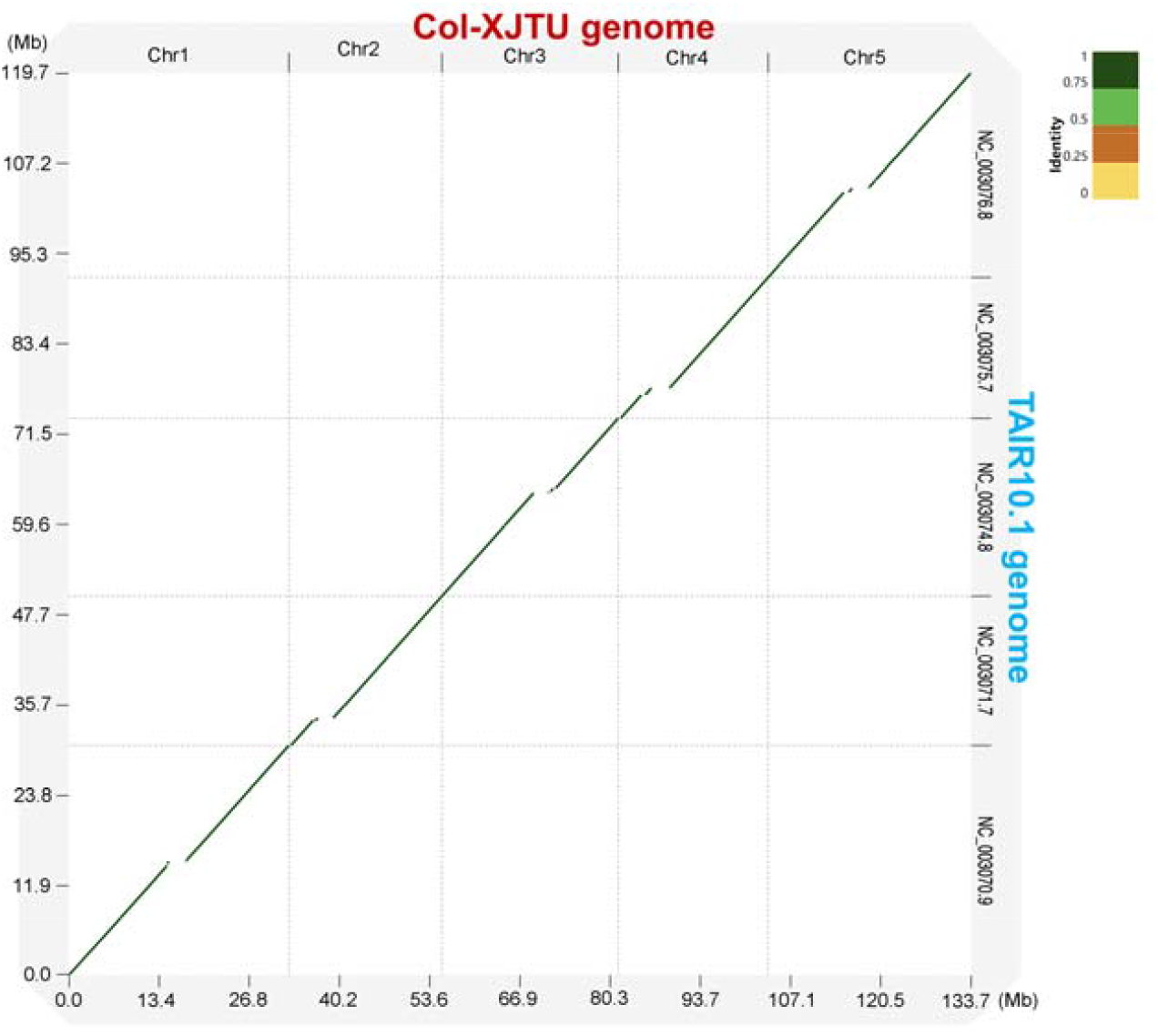
Dot plot of Col-XJTU and TAIR10.1 genome assemblies.

**Figure S4.**
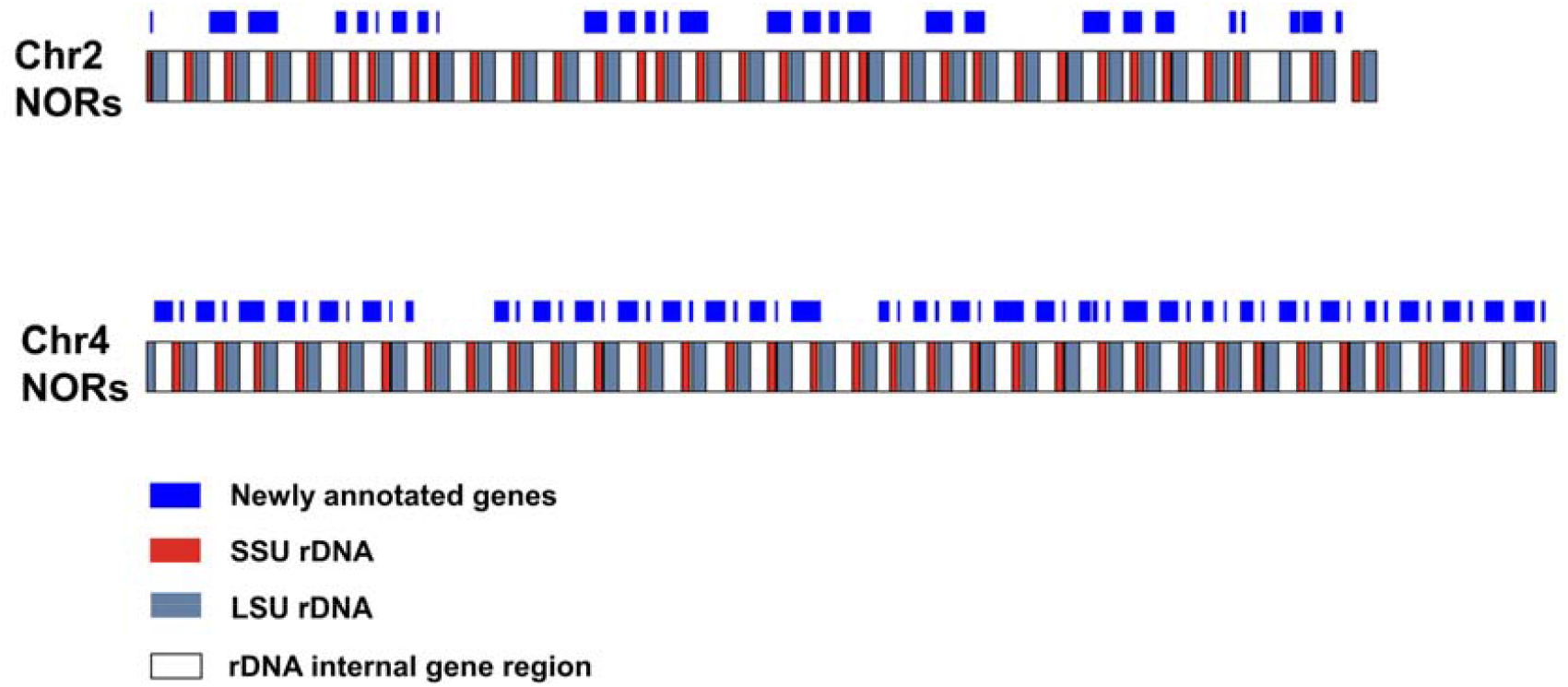
Arrangement of 45S rDNA units and new gene annotations in NORs of Chr2 and Chr4. NOR, nucleolar organizing region.

**Figure S5.**
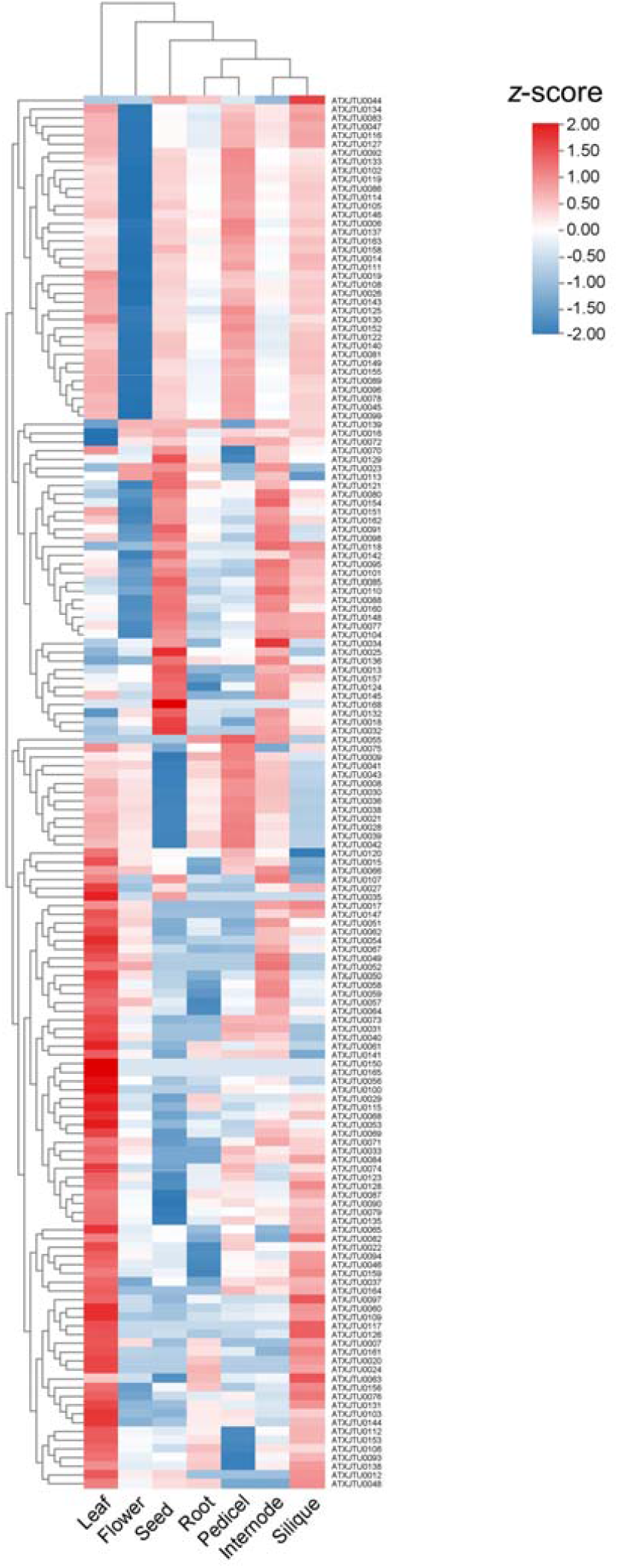
Heatmap of expression profile for newly-annotated genes in seven tissues.

**Figure S6.**
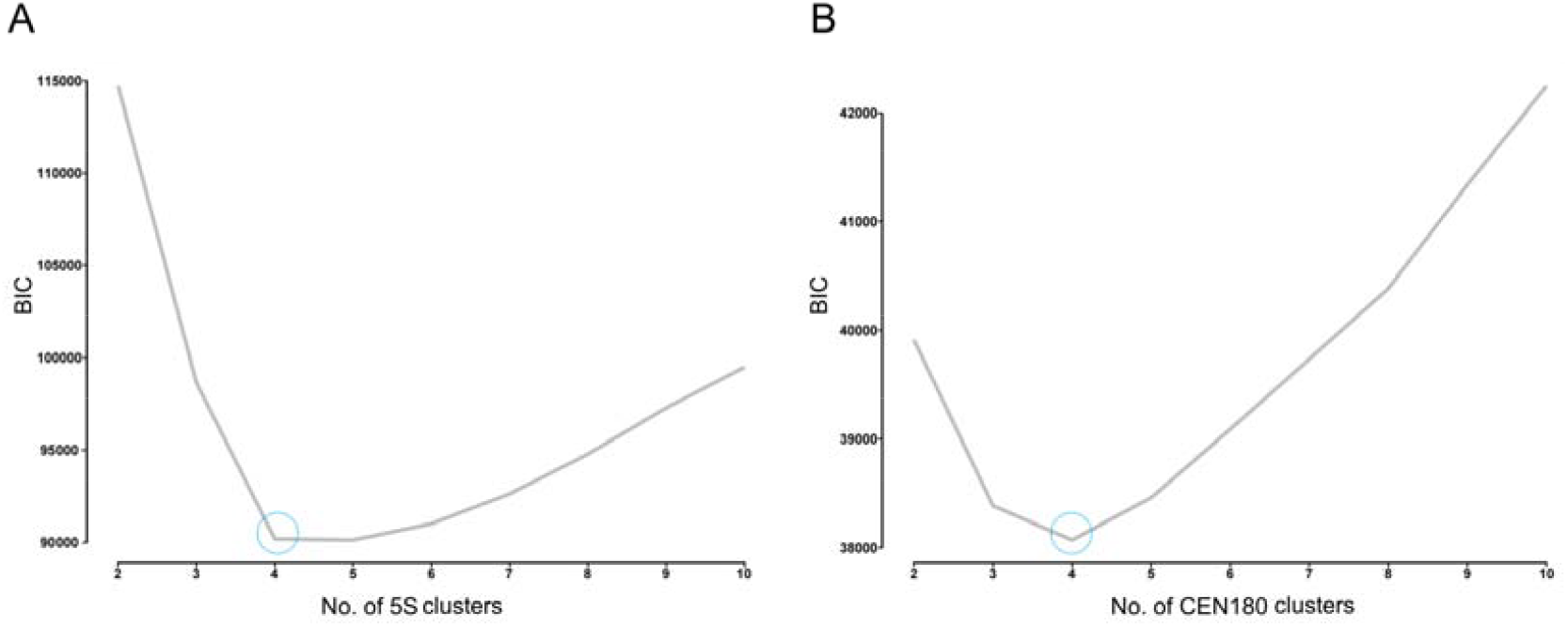
BIC inflection point plot visualized for optimal K selection. A. For 5S clusters. **B**. For CEN180 clusters.

**Figure S7.**
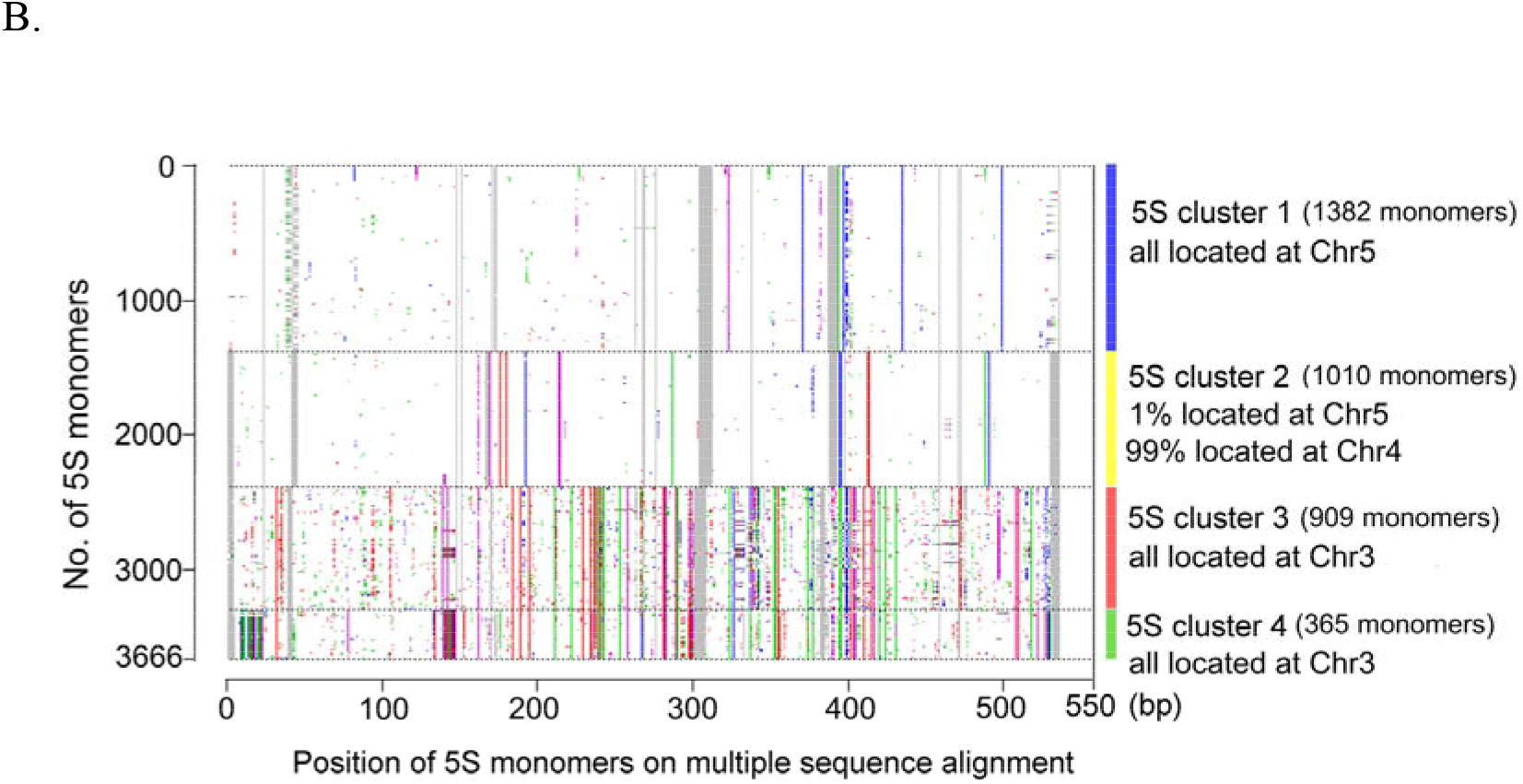
An multiple sequence alignment of the 3666 5S rDNA monomers. Segregating sites with respect to the consensus sequence are colored with green, blue, purple, red, and gray representing A, G, C, T, and a gap, respectively.

**Figure S8.**
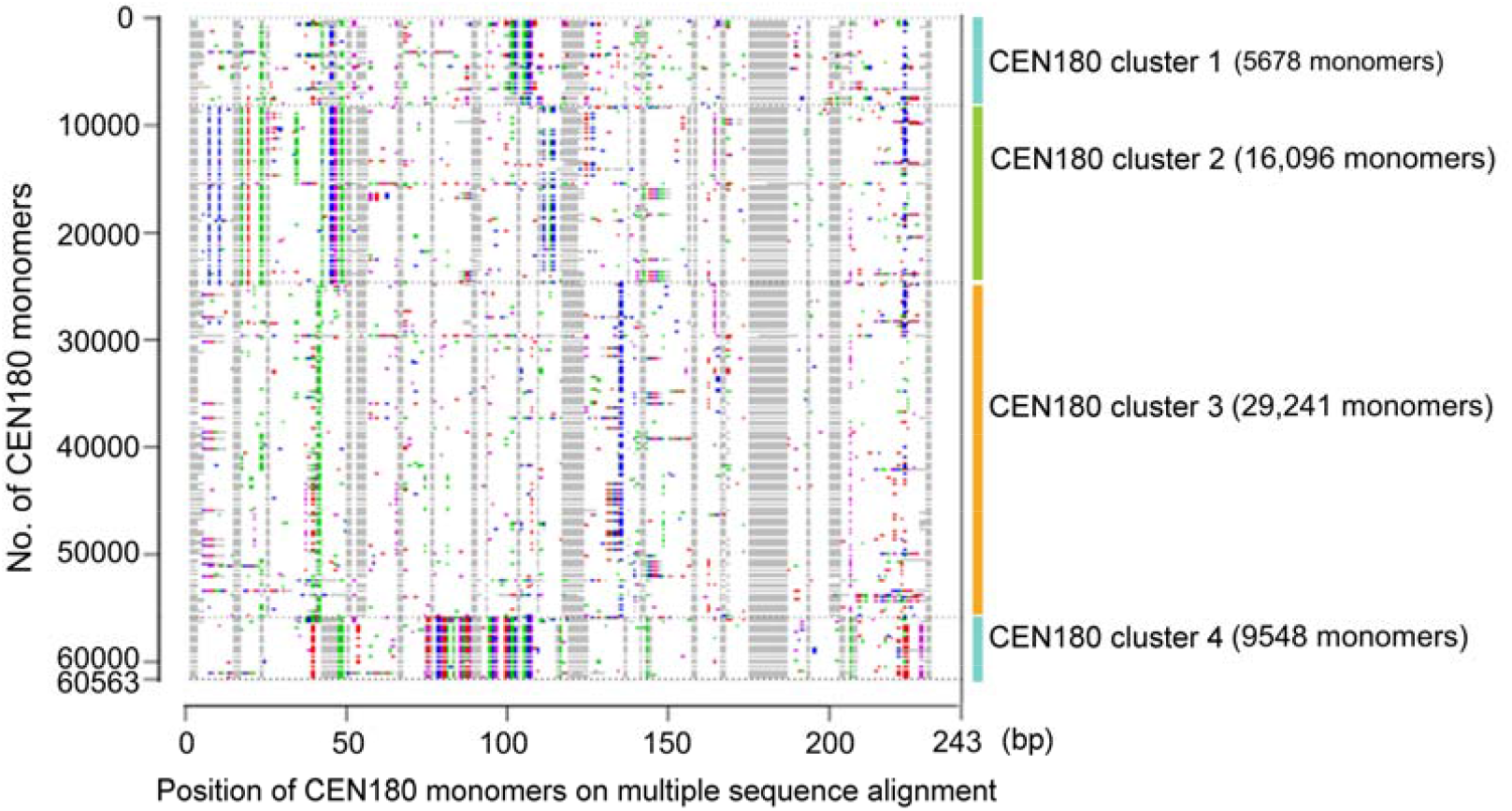
An multiple sequence alignment of the 60,563 CEN180 monomers. Segregating sites with respect to the consensus sequence are colored with green, blue, purple, red, and gray representing A, G, C, T, and a gap, respectively.

**Figure S9.**
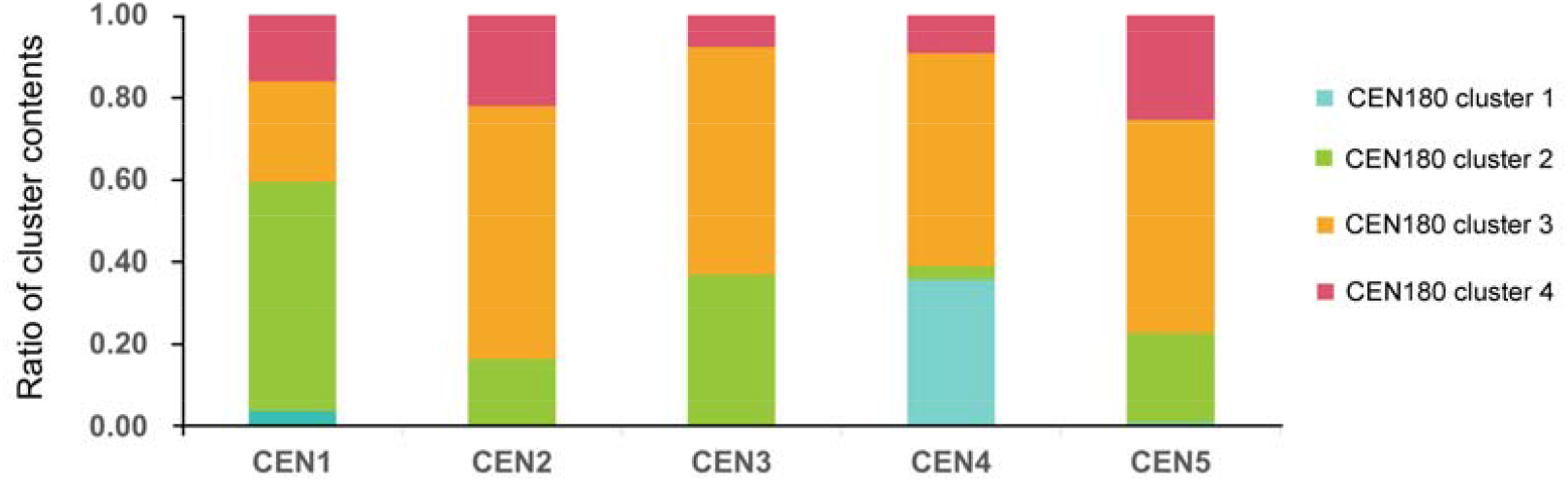
Stacked bar graph illustrating the distribution of four CEN180 clusters in five centromeres.

**Figure S10.**
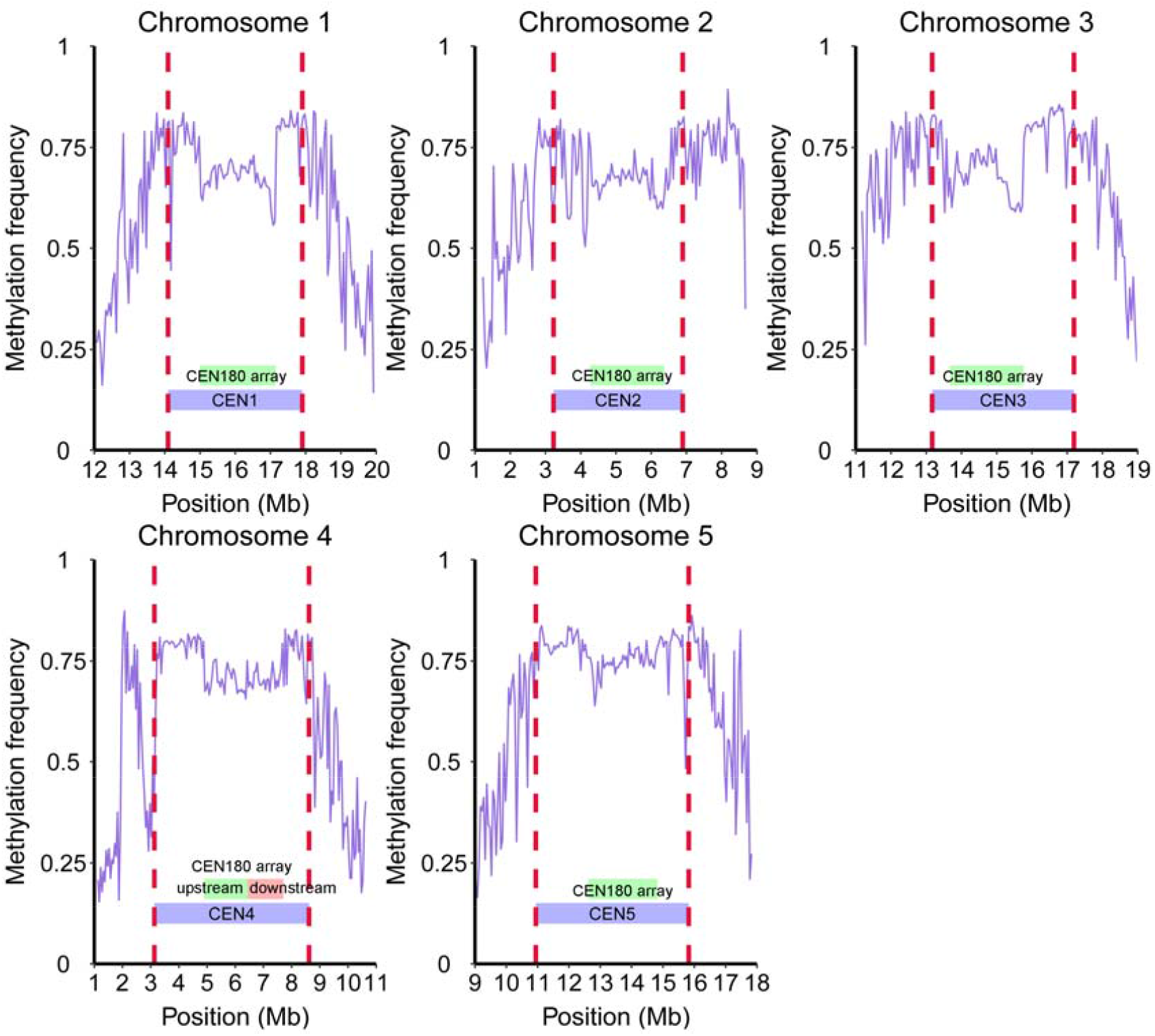
Methylation frequency in five centromeres and their pericentromeric regions (flanking 2 Mb) For each plot, two red dash lines indicates the boundaries of the respective centromere.

**Figure S11.**
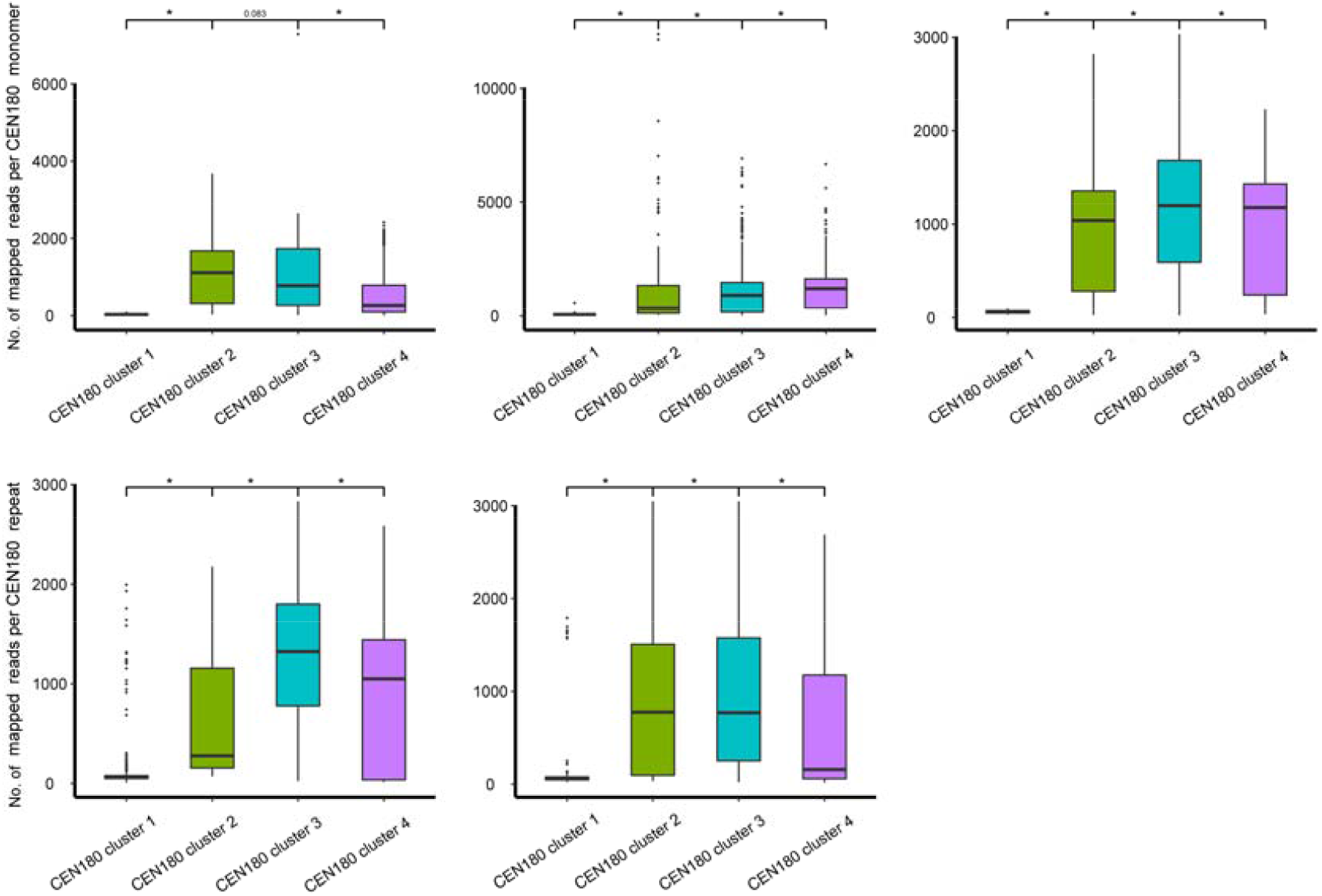
Distribution of CENH3 ChIP-seq signal across the four CEN180 clusters in five centromeres. All *P* values were calculated using Wilcoxon test. Asterisk represents significance terms (*P* < 0.05).

**Figure S12.**
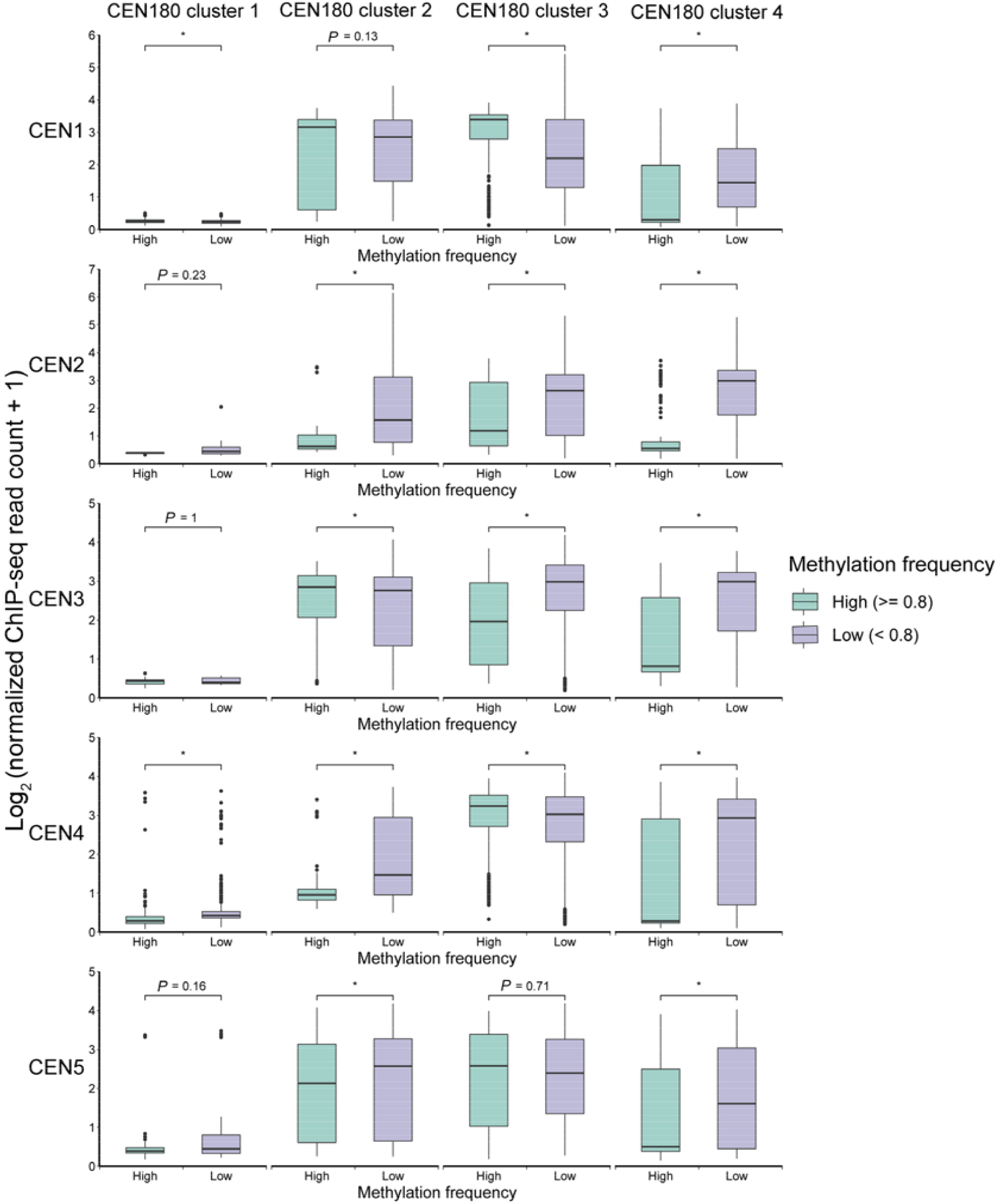
Box plots showing the methylation frequency and CENH3 binding signal across the four CEN180 clusters in five centromeres. All *P* values were calculated using Wilcoxon test. Asterisk represents significance terms (*P* < 0.05).

**Figure S13.**
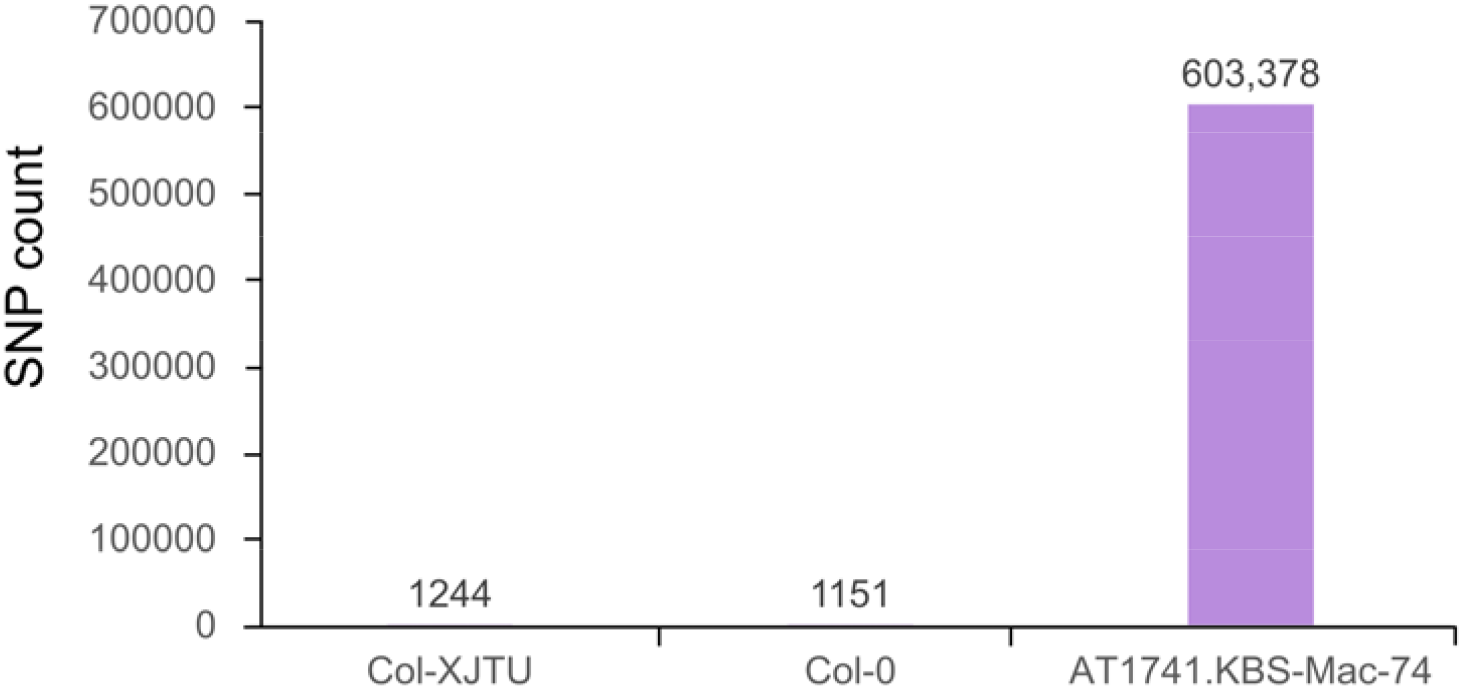
SNP count in Col-XJTU, a public wild type Col-0, and AT1741.KBS-Mac-74.

## Supplemental Tables

**Table S1 Position of ONT assembled segments in Col-XJTU assembly**

**Table S2 Misassembled regions in TAIR10.1 assembly**

**Table S3 Information of the five centromeres in the Col-XJTU assembly**

**Table S4 Gaps in the TAIR10.1 assembly**

**Table S5 Novel sequences introduced in the Col-XJTU assembly**

**Table S6 Information of Arabidopsis-type heptamer repeats CCCTAAA/TTTAGGG and 45S rDNA units in the five chromosomes**

**Table S7 Classified repeats in Col-XJTU and TARI10.1 assemblies**

**Table S8 Function description of newly-annotated genes**

**Table S9 Expression (FPKM) of newly-annotated genes in seven tissues**

**Table S10 Source of BAC sequences used as anchors and BAC validation**

