## Supplementary material for "High-quality *Arabidopsis thaliana* Genome Assembly with Nanopore and HiFi Long Reads": Table S1

**Table S1 Position of ONT assembled segments in Col-XJTU assembly**

| **Chr ID** | **Start** | **End** | **Length (bp)** | **Repeat content (%)** |
| --- | --- | --- | --- | --- |
| Chr1 | 1 | 308,852 | 308,852 | 4.66 |
| Chr1 | 23,167,339 | 23,223,725 | 56,387 | 8.40 |
| Chr2 | 2,254,183 | 3,412,890 | 1,158,708 | 54.29 |
| Chr2 | 9,719,612 | 10,875,230 | 1,155,619 | 14.18 |
| Chr2 | 15,980,222 | 16,061,133 | 80,912 | 2.32 |
| Chr2 | 22,019,922 | 22,560,461 | 540,540 | 3.60 |
| Chr3 | 13,264,628 | 13,412,425 | 147,798 | 54.41 |
| Chr4 | 4,532,360 | 4,787,126 | 254,767 | 67.39 |
| Chr5 | 3,408,744 | 3,454,912 | 46,169 | 2.12 |
| Chr5 | 15,821,034 | 16,169,952 | 348,919 | 78.50 |

*Note:* Chr, chromosome.
