## Supplementary material for "High-quality *Arabidopsis thaliana* Genome Assembly with Nanopore and HiFi Long Reads": Table S2

**Table S2 Misassembled regions in TAIR10.1 assembly**

| **Chr ID** | **Start** | **End** | **Length (bp)** | **Related protein-coding genes** |
| --- | --- | --- | --- | --- |
| NC_003076.8 | 5,775,535 | 5,776,346 | 812 | AT5G17522 |
| NC_003076.8 | 5,775,586 | 5,777,350 | 1765 | AT5G17523 |

*Note:* These regions are duplicated in Chr 5 assembled from Contig 7 (shown in green in Figure 1) in this study.
