## Supplementary material for "High-quality *Arabidopsis thaliana* Genome Assembly with Nanopore and HiFi Long Reads": Table S3

**Table S3 Information of the five centromeres in the Col-XJTU assembly**

| **Chr ID** | **Start** | **End** | **Name** | **Length (bp)** | **QV** |
| --- | --- | --- | --- | --- | --- |
| Chr1 | 14,092,143 | 17,903,680 | CEN1 | 3,811,538 | 69.91 |
| Chr2 | 3,229,032 | 6,742,512 | CEN2 | 3,513,481 | 64.00 |
| Chr3 | 13,168,960 | 17,190,023 | CEN3 | 4,021,064 | 61.78 |
| Chr4 | 3,131,982 | 8,618,139 | CEN4 | 5,486,158 | 67.90 |
| Chr5 | 10,942,051 | 15,821,029 | CEN5 | 4,878,979 | 76.43 |
