## Supplementary material for "High-quality *Arabidopsis thaliana* Genome Assembly with Nanopore and HiFi Long Reads": Table S4

**Table S4 Gaps in the TAIR10.1 assembly**

| **Gap ID** | **Chr ID** | **Start** | **End** | Length of Ns (bp) |
| --- | --- | --- | --- | --- |
| Gap1 | NC_003070.9 | 13,201,252 | 13,201,352 | 100 |
| Gap2 | NC_003070.9 | 13,993,095 | 13,993,195 | 100 |
| Gap3 | NC_003070.9 | 14,511,721 | 14,538,721 | 27,000 |
| Gap4 | NC_003070.9 | 14,545,721 | 14,581,719 | 35,998 |
| Gap5 | NC_003070.9 | 14,610,668 | 14,656,720 | 46,052 |
| Gap6 | NC_003070.9 | 14,750,970 | 14,803,970 | 53,000 |
| Gap7 | NC_003070.9 | 15,086,045 | 15,087,045 | 1000 |
| Gap8 | NC_003070.9 | 17,639,368 | 17,639,468 | 100 |
| Gap9 | NC_003070.9 | 28,537,489 | 28,537,589 | 100 |
| Gap10 | NC_003071.7 | 0 | 1000 | 1000 |
| Gap11 | NC_003071.7 | 28,030 | 28,150 | 120 |
| Gap12 | NC_003071.7 | 3,607,929 | 3,608,929 | 1000 |
| Gap13 | NC_003071.7 | 3,617,844 | 3,617,944 | 100 |
| Gap14 | NC_003074.8 | 0 | 100 | 100 |
| Gap15 | NC_003074.8 | 9,171,882 | 9,171,982 | 100 |
| Gap16 | NC_003074.8 | 11,354,159 | 11,354,259 | 100 |
| Gap17 | NC_003074.8 | 13,022,636 | 13,022,736 | 100 |
| Gap18 | NC_003074.8 | 13,587,786 | 13,588,786 | 1000 |
| Gap19 | NC_003074.8 | 13,744,449 | 13,744,549 | 100 |
| Gap20 | NC_003074.8 | 13,749,721 | 13,749,821 | 100 |
| Gap21 | NC_003074.8 | 13,799,417 | 13,800,417 | 1000 |
| Gap22 | NC_003074.8 | 14,208,952 | 14,209,952 | 1000 |
| Gap23 | NC_003074.8 | 15,132,294 | 15,132,545 | 251 |
| Gap24 | NC_003075.7 | 0 | 1000 | 1000 |
| Gap25 | NC_003075.7 | 3,053,301 | 3,054,301 | 1000 |
| Gap26 | NC_003075.7 | 3,956,021 | 3,957,021 | 1000 |
| Gap27 | NC_003076.8 | 11,194,537 | 11,194,848 | 311 |
| Gap28 | NC_003076.8 | 11,203,128 | 11,203,439 | 311 |
| Gap29 | NC_003076.8 | 11,211,026 | 11,211,337 | 311 |
| Gap30 | NC_003076.8 | 11,218,091 | 11,218,402 | 311 |
| Gap31 | NC_003076.8 | 11,223,011 | 11,223,322 | 311 |
| Gap32 | NC_003076.8 | 11,226,521 | 11,226,621 | 100 |
| Gap33 | NC_003076.8 | 11,228,286 | 11,228,386 | 100 |
| Gap34 | NC_003076.8 | 11,228,758 | 11,228,858 | 100 |
| Gap35 | NC_003076.8 | 11,725,025 | 11,726,023 | 998 |
| Gap36 | NC_003076.8 | 12,827,448 | 12,828,448 | 1000 |
| Gap37 | NC_003076.8 | 12,829,008 | 12,832,167 | 3159 |
| Gap38 | NC_003076.8 | 12,834,674 | 12,837,834 | 3160 |

*Note:* N, unidentified nucleotide.
