## Supplementary material for "High-quality *Arabidopsis thaliana* Genome Assembly with Nanopore and HiFi Long Reads": Table S6

**Table S6 Information of *Arabidopsis*-type heptamer repeats CCCTAAA/TTTAGGG and 45S rDNA units in the five chromosomes**

| **Assembly** | **Chr ID** | **5'-telomere** | | | **3'-telomere** | | |
| --- | --- | --- | --- | --- | --- | --- | --- |
|  |  | **Region** | **Sequencing length (bp)** | **Average ONT read depth (×)** | **Region** | **Sequencing length (bp)** | **Average ONT read depth (×)** |
| Col-XJTU | Chr1 | (CCCTAAA)_509_ | 3563 | 351 | (TTTAGGG)_474_ | 3318 | 456 |
|  | Chr2 | (45S rDNA unit)_33_ | 300,270 | 427 | (TTTAGGG)_378_ | 2646 | 396 |
|  | Chr3 | (CCCTAAA)_306_ | 2142 | 403 | (TTTAGGG)_433_ | 3031 | 443 |
|  | Chr4 | (45S rDNA unit)_34_ | 343,661 | 369 | (TTTAGGG)_266_ | 1862 | 444 |
|  | Chr5 | (CCCTAAA)_411_ | 2877 | 416 | (TTTAGGG)_445_ | 3115 | 473 |
| TAIR10.1 | Chr1 | (CCCTAAA)_8_ | 56 | NA | (TTTAGGG)_5_ | 35 | NA |
|  | Chr2 | (45S rDNA unit)_1_ | 5901 | NA | (TTTAGGG)_81_ | 567 | NA |
|  | Chr3 | (CCCTAAA)_9_ | 63 | NA | (TTTAGGG)_0_ | 0 | NA |
|  | Chr4 | (45S rDNA unit)_0_ | 0 | NA | (TTTAGGG)_3_ | 21 | NA |
|  | Chr5 | (CCCTAAA)_1_ | 1 | NA | (TTTAGGG)_5_ | 35 | NA |
