## Supplementary material for "High-quality *Arabidopsis thaliana* Genome Assembly with Nanopore and HiFi Long Reads": Table S7

**Table S7 Classified repeats in Col-XJTU and TARI10.1 assemblies**

| **Type** | | **Col-XJTU** | **TARI10.1** |
| --- | --- | --- | --- |
| Retroelement | | 10,209 | 3927 |
|  | SINE | 525 | 299 |
|  | LINE | 1366 | 782 |
|  | LTR element | 8318 | 2846 |
| DNA transposon | | 7480 | 4273 |
| Satellite | | 1567 | 791 |
| Simple repeat | | 36,404 | 19,032 |
| Low complexity | | 9032 | 4889 |

*Note:* All annotations were analyzed using the same pipeline described in Materials and methods section. SINE, short interspersed nuclear element; LINE, long interspersed nuclear element; LTR, long terminal repeat element.
