## Supplementary material for "High-quality *Arabidopsis thaliana* Genome Assembly with Nanopore and HiFi Long Reads": Table S9

**Table S9 Function description of newly-annotated genes**

| **Gene ID** | **Pfam** | **Domain** | **GO number** |
| --- | --- | --- | --- |
| ATXJTU0001 | PF03732 | Retrotransposon gag protein |  |
| ATXJTU0001 | PF13650 | Aspartyl protease |  |
| ATXJTU0003 | PF11690 | Protein of unknown function (DUF3287) |  |
| ATXJTU0004 | PF11690 | Protein of unknown function (DUF3287) |  |
| ATXJTU0053 | PF05405 | Mitochondrial ATP synthase B chain precursor (ATP-synt_B) | GO:0000276\|GO:0015078\|GO:0015986 |
| ATXJTU0053 | PF00420 | NADH-ubiquinone/plastoquinone oxidoreductase chain 4L |  |
| ATXJTU0056 | PF00116 | Cytochrome C oxidase subunit II, periplasmic domain | GO:0004129\|GO:0005507\|GO:0016020 |
| ATXJTU0056 | PF02790 | Cytochrome C oxidase subunit II, transmembrane domain | GO:0016021\|GO:0022900 |
| ATXJTU0059 | PF13966 | Zinc-binding in reverse transcriptase |  |
| ATXJTU0060 | PF00329 | Respiratory-chain NADH dehydrogenase, 30 kD subunit | GO:0008137\|GO:0055114 |
| ATXJTU0060 | PF00252 | Ribosomal protein L16p/L10e | GO:0003735\|GO:0005840\|GO:0006412 |
| ATXJTU0062 | PF00137 | ATP synthase subunit C | GO:0015078\|GO:0015991\|GO:0033177 |
| ATXJTU0065 | PF05919 | Mitovirus RNA-dependent RNA polymerase |  |
| ATXJTU0068 | PF00115 | Cytochrome C and quinol oxidase polypeptide I | GO:0004129\|GO:0009060\|GO:0016021\|GO:0020037\|GO:0055114 |
| ATXJTU0071 | PF00146 | NADH dehydrogenase | GO:0016020\|GO:0055114 |
| ATXJTU0073 | PF00146 | NADH dehydrogenase | GO:0016020\|GO:0055114 |
| ATXJTU0074 | PF18044 | CCCH-type zinc finger |  |
| ATXJTU0074 | PF18044 | CCCH-type zinc finger |  |
| ATXJTU0074 | PF00097 | Zinc finger, C3HC4 type (RING finger) | GO:0046872 |
| ATXJTU0165 | PF03078 | ATHILA ORF-1 family |  |
