## Supplementary material for "High-quality *Arabidopsis thaliana* Genome Assembly with Nanopore and HiFi Long Reads": Table S10

**Table S10 Source of BAC sequences used as anchors and BAC validation**

| **BAC ID used as anchor** | **GenBank accession No.** | **Chr ID** | **Length (bp)** | **Link for BAC sequence of Chr used for BAC validation** |
| --- | --- | --- | --- | --- |
| F22M8 | AC020622.3 | Chr1 | 47,295 | <https://www.ebi.ac.uk/genomes/ath1.html> |
| F14J16 | AC002304.3 | Chr1 | 106,320 |  |
| F14G9 | AC069159.8 | Chr1 | 111,481 |  |
| F5G3 | AC007018.7 | Chr2 | 124,326 | <https://www.ebi.ac.uk/genomes/ath2.html> |
| F27C21 | AC006527.5 | Chr2 | 33,117 |  |
| T12J2 | AC004483.3 | Chr2 | 85,463 |  |
| T15D9 | AC007120.6 | Chr2 | 102,331 |  |
| F9O13 | AC006248.4 | Chr2 | 119,753 |  |
| F24H14 | AC006135.3 | Chr2 | 82,356 |  |
| T11J7 | AC002340.3 | Chr2 | 79,574 |  |
| T16B12 | AC005311.3 | Chr2 | 82,594 |  |
| F13A10 | AC006418.4 | Chr2 | 84,825 |  |
| T25N22 | AC005693.3 | Chr2 | 80,891 |  |
| T5E7 | AC006225.3 | Chr2 | 88,249 |  |
| F6H5 | AP002035.1 | Chr3 | 95,681 | <https://www.ebi.ac.uk/genomes/BA000014.html> |
| T7B9 | AP002067.2 | Chr3 | 84,711 |  |
| T14K23 | AL132909.1 | Chr3 | 101,786 |  |
| T1J24 | AF147263.1 | Chr4 | 122,529 | <https://www.ebi.ac.uk/genomes/AJ270060.html> (long arm); <https://www.ebi.ac.uk/genomes/AJ270058.html> (short arm) |
| T24C1 | AB073158.1 | Chr4 | 90,840 |  |
| T7K6 | AB073163.1 | Chr4 | 31,230 |  |
| T17A2 | AF160183.1 | Chr4 | 81,902 |  |
| MAJ23 | AL392144.1 | Chr5 | 42,694 | <https://www.ebi.ac.uk/genomes/BA000015.html> |
| T5K6 | AL391222.1 | Chr5 | 96,379 |  |
| F3F24 | AC018632.1 | Chr5 | 151,828 |  |
| T15F17 | AF262042.1 | Chr5 | 68,352 |  |
| F18A12 | AC069553.6 | Chr5 | 133,529 |  |
| T5E15 | AC019013.2 | Chr5 | 114,056 |  |
| K3M16 | AL391150.1 | Chr5 | 26,604 |  |
| K10A8 | AL391151.1 | Chr5 | 43,387 |  |
| F24C7 | AP002029.1 | Chr5 | 73,663 |  |
